# Color morphs of the coral, *Acropora tenuis*, show different responses to environmental stress and different expression profiles of fluorescent-protein genes

**DOI:** 10.1101/2020.08.28.272823

**Authors:** Noriyuki Satoh, Koji Kinjo, Kohei Shintaku, Daisuke Kezuka, Hiroo Ishimori, Atsushi Yokokura, Kazutaka Hagiwara, Kanako Hisata, Mayumi Kawamitsu, Koji Koizumi, Chuya Shinzato, Yuna Zayasu

## Abstract

Corals of the family Acroporidae are key structural components of reefs that support the most diverse marine ecosystems. Due to increasing anthropogenic stresses, coral reefs are in decline. Along the coast of Okinawa, Japan, three different color morphs of *Acropora tenuis* have been recognized for decades. These include brown (N morph), yellow-green (G) and purple (P) forms. The tips of axial coral polyps exhibit specific fluorescence spectra. This attribute is inherited asexually, and color morphs do not change seasonally. In Okinawa Prefecture, during the summer of 2017, the N and P morphs experienced bleaching, in which some N morphs died while P morphs recovered. In contrast, G morphs successfully withstood the stress. Symbiotic dinoflagellates are essential symbiotic partners of scleractinian corals. Photosynthetic activity of symbionts was reduced in July in N and P morphs; however, the three color-morphs host similar sets of Clade-C zoothanthellae, suggesting that beaching of N and P morphs cannot be attributed to differences in symbiont clades. The decoded *Acropora tenuis* genome includes five genes for green fluorescent proteins (GFP), two for cyan fluorescent proteins (CFP), three for red fluorescent proteins (RFP), and seven genes for chromoprotein (ChrP). A summer survey of gene expression profiles demonstrated that (a) expression of CFP and REP was quite low in all three morphs, (b) P morphs expressed higher levels of ChrP, (c) both N and G morphs expressed GFP highly, and (d) GFP expression was reduced in N morphs, compared to G morphs, which maintained higher levels of GFP expression throughout the summer. Although further studies are required to understand the biological significance of these color morphs of *Acropora tenuis*, our results suggest that thermal stress resistance is modified by genetic mechanisms that coincidentally lead to diversification of color morphs.

---

One of the most critical issues facing the human race is warming of our planet. Anthropogenic activities have harmed the environment in various ways, and coral reefs have been especially impacted (Hoegh-Guldberg *et al*. 2007; Hughes *et al*. 2017). In spite of the fact that coral reefs occupy only 0.2% of the ocean area, they are estimated to harbor about one-third of all described marine species (Knowlton *et al*. 2010; Fisher *et al*. 2015). Coral reefs are the most diverse marine ecosystems on Earth (Wilkinson 2008). Scleractinian corals, a keystone component of calcium-carbonate based reefs, form obligate endosymbiosis with photosynthetic dinoflagellates or zoothanthellae, which supply the vast majority of their photosynthetic products to the host corals (Yellowlees *et al*. 2008). However, corals now face a variety of environmental stresses, including increasing surface seawater temperatures, decimation by outbreaks of crown-of-thorns starfish, and acidification and pollution of oceans (Hoegh-Guldberg *et al*. 2007; Uthicke *et al*. 2009; Burke *et al*. 2011; Uthicke *et al*. 2015; Hughes *et al*. 2017). Loss of coral reefs is potentially catastrophic.

Faced with such environmental changes, not only are corals diminished by bleaching and other causes, but to survive they must adapt somehow to increasingly stressful environments (Skelly *et al*. 2007). *Acropora tenuis* is one of the most important scleractinian corals along the coast of Okinawa, Japan (Omori *et al*. 2016). Most *A. tenuis* appear brownish (Fig. 1a, b), reflecting the color of the symbiotic algae that they host. *Acropora tenuis* exhibits faster growth than other *Acropora* species, forming colonies approximately 30 cm in diameter within 3∼5 years (Fig. 1f) (Iwao *et al*. 2010). In 1998, along the Okinawa coast, various *Acropora* species suffered extensive bleaching and many died (Reef Conservation Committee of Japanese Coral Reef Society, 2014). After several years, they gradually recovered. However, while color morphs of *Acropora tenuis* have been described (Nishihira and Veron 1995), divers in Okinawa noticed the re-appearance of greenish-yellow and purple *A. tenuis* morphs (Fig. 1c, d) (Fig. 1e).

**Figure 1.**
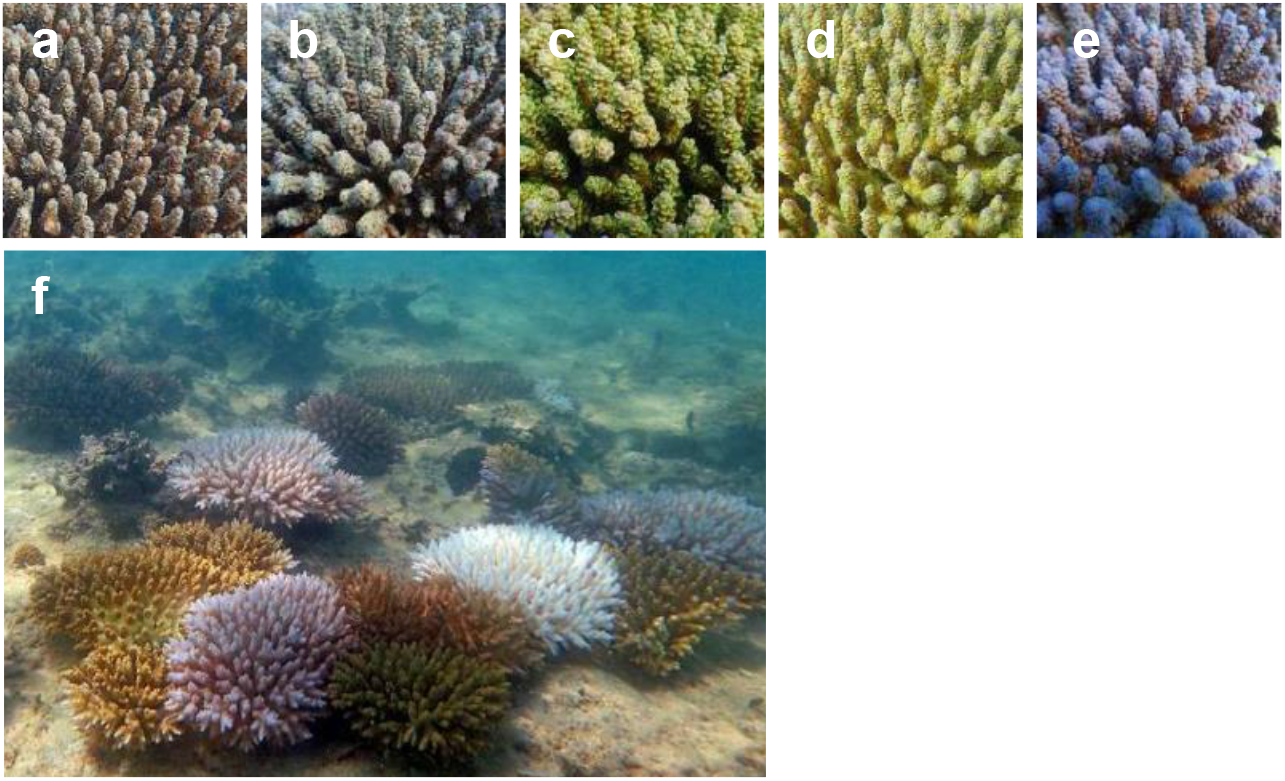
Color morphs of *Acropora tenuis* along the Okinawa coast. (a, b) Two “wild-type” morphs, NO (a) and NB (b). Both appear generally brownish while the tips of axial polyps are orange in NO (a) and blue in NB (b). (c, d) Two greenish morphs, GO (c) and GB (d). Tips of axial polyps are orange in GO (c) and blue in GB (d). (e) A purple colony (P). (f) Branches originating from color morphs are growing in shallow water off the coast of Okinawa.

In corals, fluorescent protein-mediated color polymorphism is thought to contribute to their acclimatization potential (Dove *et al*. 2001; Dove 2004; Kelmanson and Matz 2003; Salih *et al*. 2010; Smith *et al*. 2013; Paley 2014; Gittins *et al*. 2015; Jarret *et al*. 2017). Therefore, we thought that the color polymorphism of *Acropora tenuis* might be associated with its potential to resist to stressful environments. We describe here the characteristics of the three color-morphs and their capacity to accommodate rising summer surface seawater temperatures. Since the genome of *Acropora tenuis* has been decoded (Shinzato *et al*. 2020), we surveyed genes for fluorescent proteins to document the presence of genes for green fluorescent protein (GFP), red fluorescent protein (RFP), cyan fluorescent protein (CFP) and chromoprotein (ChrP). We examined gene expression profiles in the three color morphs in response to rising summer surface seawater temperatures.

## MATERIALS AND METHODS

### *Acropora tenuis* colonies with different color morphs

Various *Acropora tenuis* color morphs exist along the Okinawa coast. The “wild-type” color morph (N) looks brownish, presumably reflecting the color of its symbiotic algae. Depending upon the color of axial polyp tips, they are further subdivided into NO (the morph is brownish overall, but the tips of axial polyps are orange; Fig 1a) and NB (a brownish morph with bluish axial polyp tips; Figs. 1b and 2a). The color morph that looks greenish (G) is also subdivided into GO (axial polyp tips look orange; Fig. 1c) and GB (axial polyp tips look bluish; Figs. 1d and 2b). Yet another color morph is purple (Figs. 1e and 2c). The five morphs were collected in about 2005/2006 and have been maintained in the “Sea Seed Aquarium” (a non-governmental organization founded by K. Kinjo) at Yomitan, Okinawa, Japan.

**Figure 2.**
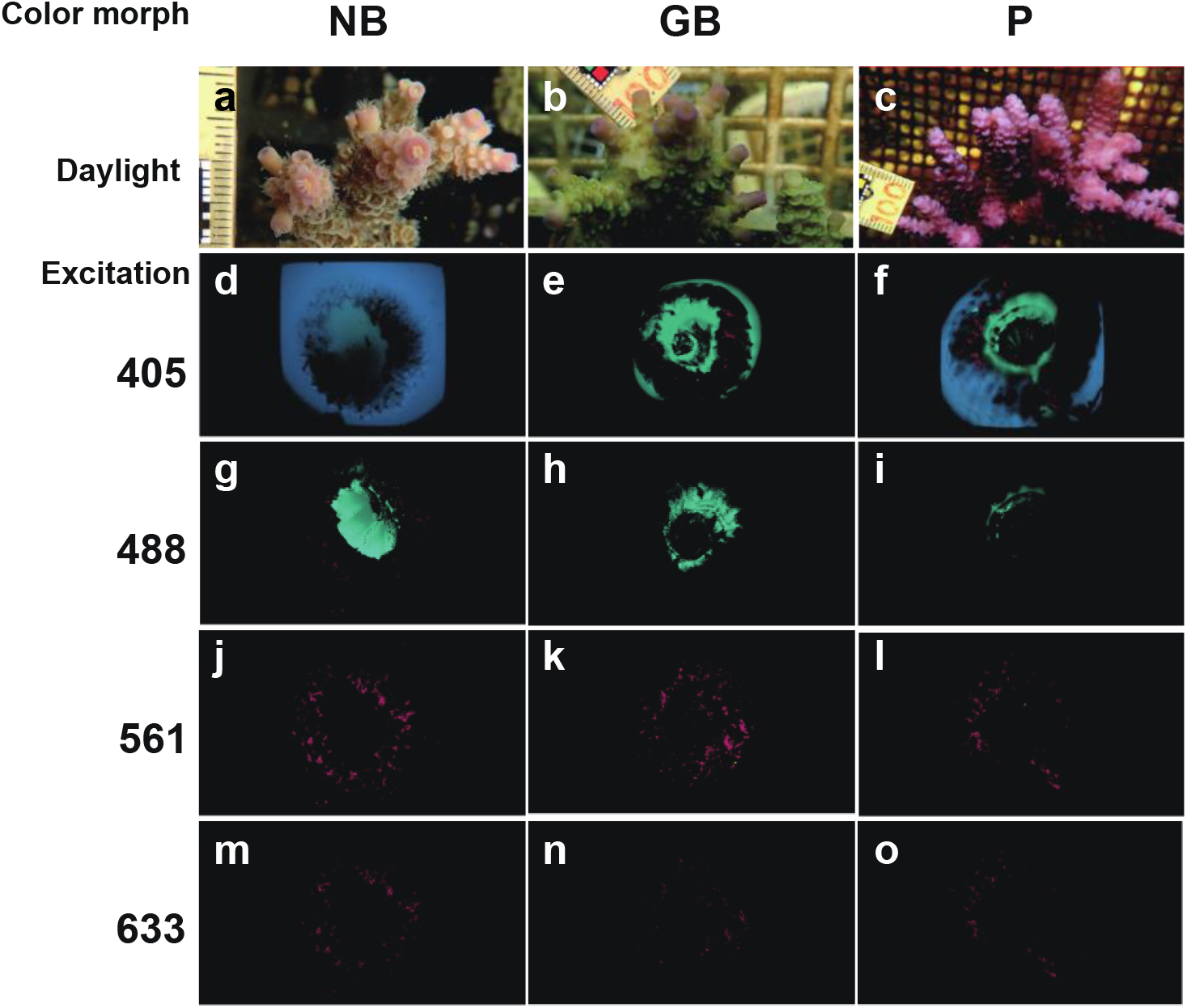
Spectral imaging of fluorescent emission at the tips of axial bodies of NB, GB, and P morphs in *A. tenuis*. Morphology of tip regions of the axial bodies of NB (a), GB (b) and P (c) color morphs. Lasers of four different wavelengths, 405 nm (d-f), 488 nm (g-i), 561 nm (j-l) and 633 nm (m-o), were used for excitation. Spectra data are displayed with wavelength color codes. Three-dimensional images were constructed with ImageJ.

Branches of wild specimens of the five different color morphs were broken off and raised for several years in the Sea Seed Aquarium. Nursery-raised corals more than 3 cm in diameter were then transplanted back into the ocean by attaching them to rocks along the shallow coast (2∼4 m in depth) in the South China Sea. After several years, they became mature colonies more than 20 cm in diameter (Fig. 1f), which were used for field observations mentioned below.

For molecular biological analyses, three to five colonies of each color morph were cultured in outdoor pools of the Sea Seed Aquarium under continuous natural seawater supply. Previously, we developed a microsatellite genotyping method that can be widely applied to *Acropora* species, including *A. tenuis*, to identify individual colony by assessing specific heterogeneity (Shinzato *et al*. 2014). This method was employed to determine genetic relationships among the five color-morphs used in this study. Each morph had an independent genotype, and genotypes of the color morphs reproduced asexually were identical (data not shown). That is, it is unlikely that color polymorphism arose from a single founder or a small group of founders. Rather, it appears that each morph was derived from widespread diversification of different ancestral morphs.

Two or three branches approximately 2 cm in length were cut from NB, GB, GO and P color-morphs on June 9th, July 26th, August 15^th^, and September 14^th^, 2017. Branches were immediately fixed in RNAlater in 50-mL Falcon tubes and were maintained at - 80°C until use. Temperature changes of the outdoor pools were recorded with a HOBO Pendant Temperature logger (Onset) from July 27th to September 27^th^. Temperatures of the pools were kept slightly lower than those of the ocean itself.

### Fluorescent Emissions

Fluorescence spectra of axial polyp tips of *A. tenuis* color morphs were examined using a Carl Zeiss LSM780 inverted confocal microscope with a 5x Fluar lens. Live branches of NB, GB and P colonies were embedded in a 1% agarose gel in a glass-bottomed dish. Lasers of four wavelengths (405 nm, 488 nm, 561 nm and 633 nm) were employed. Z-stack images, including spectral information from 409 – 695 nm were acquired using 32-channel detectors. Spectra data were displayed with wavelength color codes. Three-dimensional images were constructed with ImageJ (http://imagej.nih.gov/ij/).

### Field observations

Field observations were conducted during the summers of 2016 and 2017. An ortho-image of the research field constructed with Agisoft Metashape is shown in Fig. S1. Surface seawater temperatures at approximately 2 m depth were automatically recorded with a Compact CT (Fig. S2a) (JFE Advantech Co., LTD, Nishinomiya, Japan). The photosynthetic photon flux density (PPFD) was also recoded automatically using a DEFI2-L (Fig. S2b) (JFE Advantech Co., LTD). In 2017, changes in temperature and insolation were observed for 171 days from June 11 to November 28.

Field survey sites employed 2m X 2m quadrats (Fig. S1). Bleaching status of 169 colonies (125 NO, 35 NB, 3 GO, 3 GB, and 3 P) was determined by direct observation by divers. The degree of bleaching was quantified as the bleached surface area relative to the total surface area of each morph. Bleaching outcome (death or survival) was also recorded.

### Photosynthetic activity

Photosynthetic activity of *Acropora tenuis* colonies of different color morphs was continuously monitored at each field site. Chlorophyll fluorescence measurements were performed with a pulse amplitude-modulated fluorometer, Diving-PAM (Heinz Walz Company, Germany). Photosynthetic activity was measured by maximum quantum yield of PSII, determined as Fv/Fm. Fv was obtained as Fm-Fo, where Fo is the minimum fluorescence obtained under the measuring beam of PAM (weak pulsed light < 1μmol photons m^-2^ s^-1^) in dark-adapted condition, and Fm is the maximum fluorescence detected using a short, saturating pulse. Data were analyzed with ANOVA using JMP software (https://www.jmp.com). Statistical significance of differences was tested with the Steel-Dwass method.

### Identification of symbiotic dinoflagellates

Symbiotic dinoflagellate clades from each color morph were identified using RNA-seq data, following the method of Shinzato *et al*. (2018). Briefly, quality trimmed (QV>20) sequencing reads were aligned to the curated database of Symbiodiniaceae internal transcribed spacer 2 from NCBI using BLASTN (Camacho *et al*. 2009) with an e-value cutoff of 1e^-20^, and the number of reads that ensured that the BLASTN bit score of the best alignment was greater than the second-best alignment was counted.

### The *Acropora tenuis* genome, isolation of genes for fluorescent proteins, and molecular phylogeny

A draft genome of *Acropora tenuis* was sequenced using Illumina technology by Shinzato *et al*. (2020). The approximately 407-Mbp *A. tenuis* genome is estimated to contain 23,119 protein-coding genes with a BUSCO score of 97.4% (92.2% completeness). *Acropora tenuis* genes for green (GFP), cyan (CFP), and red (RFP) fluorescent proteins, and non-fluorescent blue/purple chromoprotein (ChrP) were identified by BLAST searching of the genome using *A. digitifera* genes as queries for GFP, CFP, RFP and ChrP, respectively (Shinzato *et al*. 2012). Orthologous relationships of genes of the two *Acropora* species were confirmed by molecular phylogenetic analysis, carried out using RAxML 8.2.11 (Stamatakis 2016)

### Expression levels of genes for GFP, CRP, RFP and ChrP

Total RNA was extracted from all specimens using an RNA plant mini Kit (Qiagen). cDNA libraries were produced using a TruSeq Standard mRNA Library Prep Kit for NeoPrep (Illumina). Library sequencing was carried out on a Hi-seq4000 (Illumina) according to the method described by Pertea *et al*. (2015). Sequencing adapters and low-quality (<Q30) sequence of Illumina reads were trimmed with Trimmomatic 0.33 (Bolger *et al*. 2014) and PCR duplicate sequences were removed with prinseq 0.20.3 (Schmieder and Edwards 2011). These trimmed sequence data were mapped onto the genome with HISAT2 2.1.0 (Pertea *et al*. 2014) and counted with featureCounts 1.6.5 (Liao *et al*. 2014). Expression degree analysis was performed using edgeR 3.28.1 (McCarthy *et al*. 2012) and DESeq2 1.26.0 (Love *et al*. 2014).

## RESULTS

### Color morphs of *Acropora tenuis* at the Okinawa coast

The Ryukyu Archipelago, and especially the Okinawa coast, experienced very severe coral bleaching in 1998. Afterward, local divers and researchers noticed that recovering *A. tenuis* exhibited different color morphs (Fig. 1). These included green (GO and GB) and purple morphs (P), as well as the predominant brown (NO and NB) color morphs (Fig. 1). These were collected and maintained at the Sea Seed Aquarium. Nubbins of each color morph were asexually reproduced by cutting branches from original colonies, nursing them for a few years at the Aquarium, and culturing them in the ocean (average depth, 2∼4 m) adjacent to the Aquarium (Fig. 1f). Since then, the five color morphs have maintained their originals colors and features.

To characterize the color morphs, fluorescence spectral imaging was carried out on the tips of axial polyps of NB, GB, and P morphs. Lasers (405, 488, 561, and 633 nm) were employed, and three-dimensional images were constructed using ImageJ. Polyps of all color morphs fluoresced green inside the mouth when excited at 488 nm (Fig. 2g-i). On the other hand, the response to 405-nm excitation differed among color morphs. Body walls of NB and P morphs fluoresced blue (Fig. 2d, f). In contrast, body walls of GB morphs fluoresced green (Fig. 2e). Fewer differences in red and deep red fluorescent emissions were detected among the three color morphs when excited at 561 and 633 nm (Fig. 2j-o). Although detailed molecular and biochemical mechanisms remain to be explored, these differences in fluorescence emission may be associated with differences in the color of *A. tenuis*.

### Responses of color morphs to summer environmental stress

To examine whether the three color morphs differ in their potential to respond to environmental changes, especially the rise of surface seawater temperatures, we followed bleaching profiles of the five morphs in the summers of 2016 and 2017. A similar trend in the mode of bleaching was observed in both years. Since the seawater temperature rise was higher and coral bleaching was more severe in 2017 than 2016, we describe here results from 2017.

Changes in surface seawater temperatures (Fig. S2a) and photosynthetic photon flux density (PPFD) (Fig. S2b) were automatically recorded using a Compact CT and a DEFI2-L, respectively. The surface seawater temperatures rose to over 30°C in early July, a temperature that continued until early September. Forty-six of 171 days exceeded 30°C, with a maximum of 34.2°C on July 27 (Fig. S2a). The PPFD also increased toward mid-July, and a high level of PPFD continued until late September, although daily variations occurred (Fig. S2b). The highest was 4,187 μ mol/m^-2^ sec^-1^ on August 25. Bleaching status was observed by diving on July 26, August 30, September 29, and November 29.

The degree of bleaching was highest in NO and NB, then P (Fig. 3). More than half of NO colonies commenced bleaching in July and August (Fig. 3a). Although 20∼30% of bleached NOs recovered their color morphs in September and November, 10∼15% of them died after bleaching (Fig. 3a). The percentage of bleached NBs in July and August was less than that of NOs, but many of the bleached NBs did not recover and died (Fig. 3b). Nearly 30% of NBs died by late November (Fig. 3b). Nearly 20% of the P morph surface area appeared dead (Fig. 3e). Bleaching occurred in 30% of P morphs in late August, but most of them recovered in late September (Fig. 3e). In contrast, G morphs, especially GB, did not bleach at all (Fig. 3d), although a few GO colonies died for unknown reasons (Fig. 3c). This suggests that G morphs have the greatest potential resistance to increasing summer surface seawater temperatures in Okinawa *A. tenuis*.

**Figure 3.**
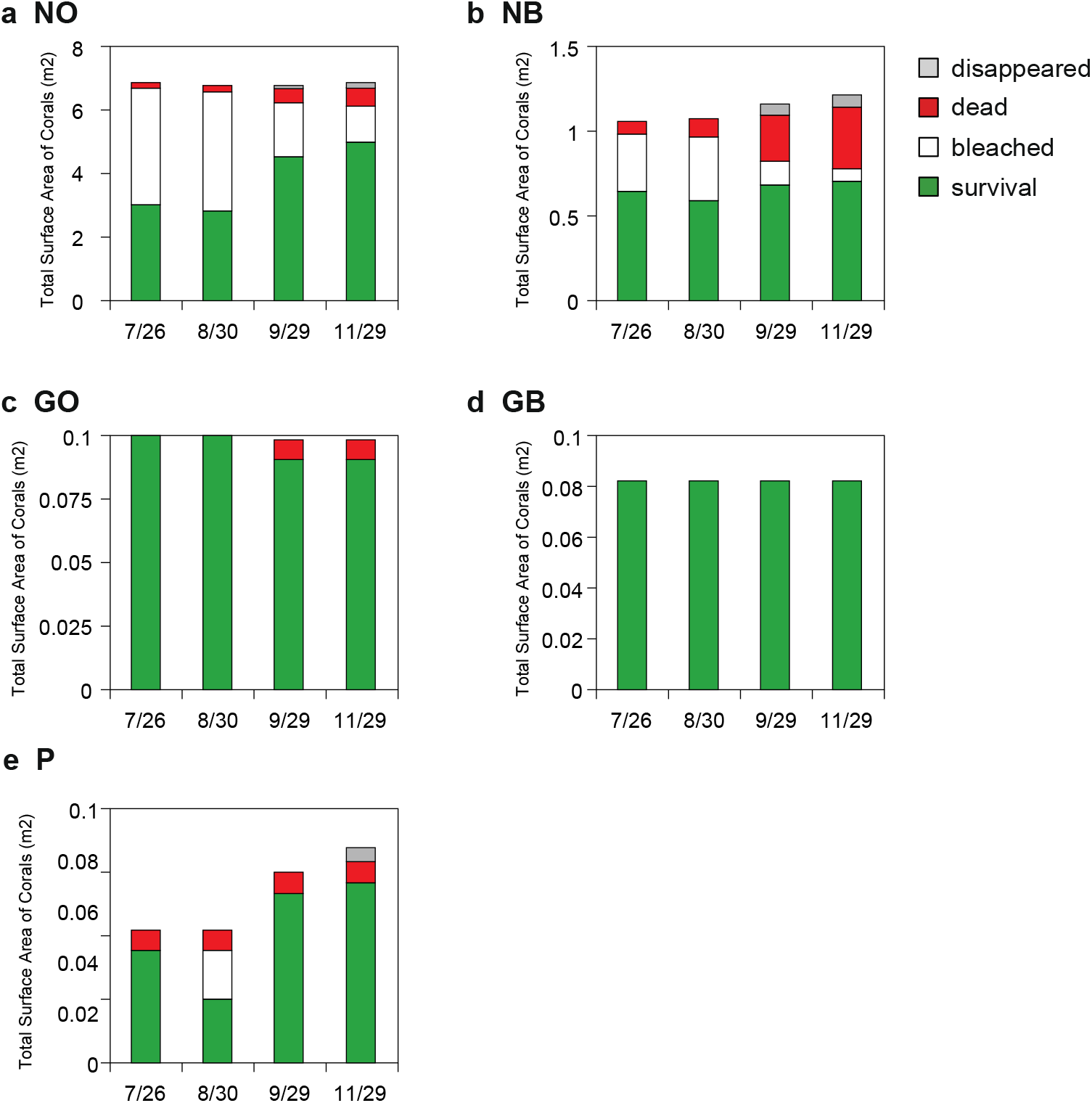
Bleaching and subsequent death of *Acropora tenuis* color morphs in the summer of 2017 along the Yomitan, Okinawa coast. The surface area of color morphs that exhibit bleaching is shown in white, with that of healthy areas is green. Dead areas after bleaching are in red. Colonies that disappeared due to wave action or unknown causes are shown in grey. Because the numbers of each color morph examined differ, areas are shown by different values on the y-axis (m^2^). It is evident that GO and GB were little affected by environmental stresses, compared to NO, NB and P.

Coral bleaching is caused by the escape of symbiotic dinoflagellates from corals or by their death inside corals. Decreased photosynthetic activity of symbiotic dinoflagellates may degrade their physical condition, and in turn, that of coral hosts. Photosynthetic activity of symbionts can be measured by chlorophyll fluorescence (Fv/Fm). We examined whether the three color-morphs show different Fv/Fm profiles (Fig. 4). The observed Fv/Fm value was nearly the same in all five color morphs in June, August, September and November, but in July, GB and GO maintained their levels of Fv/Fm, whereas it was reduced in NO, and was lowest in P and NB (Fig. 4a). GB and GO maintained photosynthetic activity at the level of June (Fig. 4a). While activity was reduced in P and NB, both recovered their photosynthetic rates in August to the same level as in GB and GO (Fig. 4a). The decrease in Fv/Fm of NB and P in July was statistically significant, compared to those of GB and GO (p<0.05) (Fig. 4b). The positive correlation between the rate of bleaching and photosynthetic activity confirms that coral bleaching is caused at least to some extent by decreased photosynthetic activity of symbiotic dinoflagellates, as shown by previous studies (e.g., Smith *et al*. 2013; Gittins *et al*. 2015).

**Figure 4.**
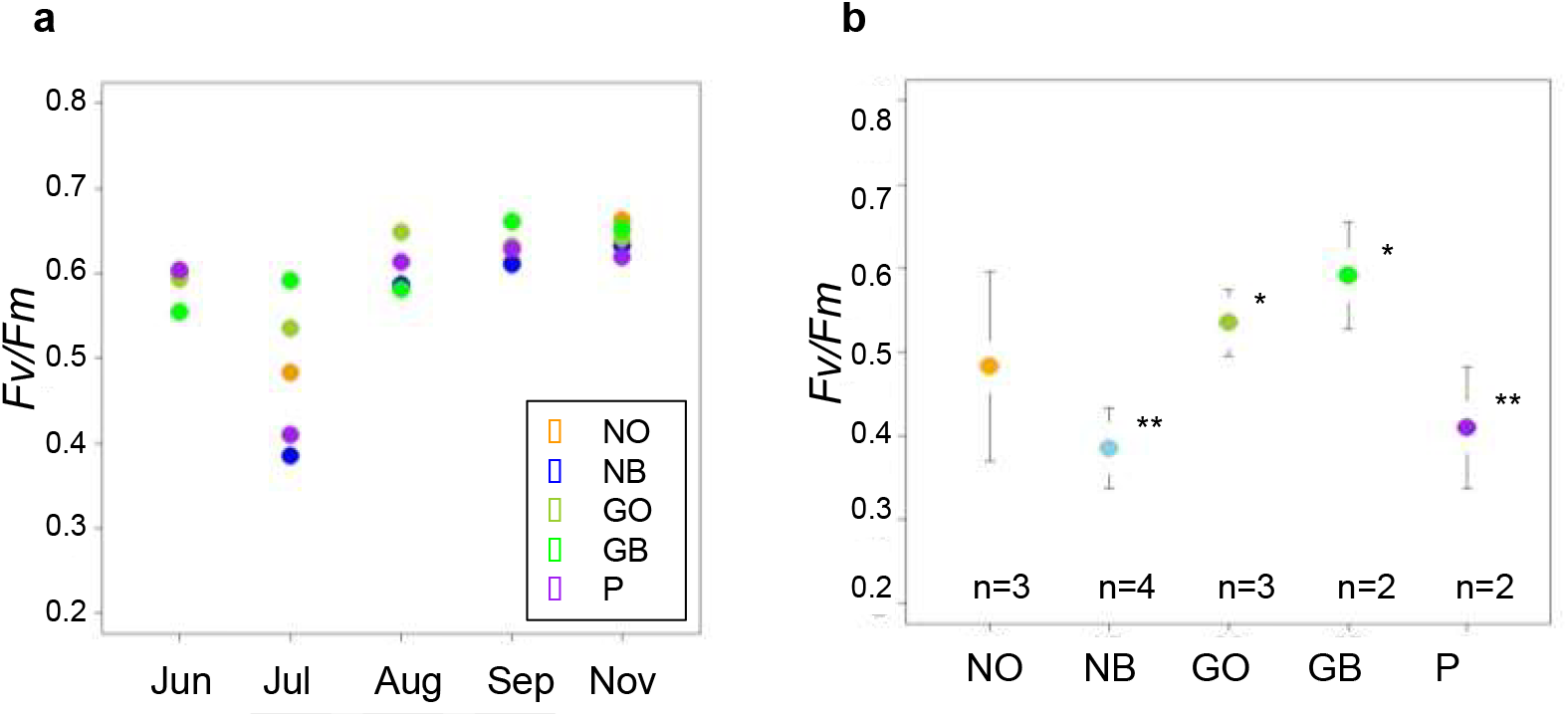
Photosynthetic activity (*Fv/Fm*) of five color morphs of *Acropora tenuis* during the summer of 2017. (a) Activity profiles in June, July, August, September and November. Differences in activity were evident in July among the five color morphs. (b) Comparison of activities in July among the five color morphs. n, the number of colonies examined. Bars indicate one standard deviation. * and ** show statistically-relevant differences between GO/GB and NB/P, using the Steel-Dwass method.

### Clade distribution of symbiotic dinoflagellates among color morphs

One possible explanation for the differing potentials of the various color morphs to resist thermal stress is that they host different clades of symbiotic dinoflagellates. The zoothanthella family Symbiodiniaceae has recently been reorganized into nine clades or genera, according to new analyses of combinatorial data (LaJeunesse *et al*. 2018). It has been suggested that corals that host some species of Clade D (*Durusdinium*) are more heat-resistant than those that host species of Clade C (*Cladocopium*). Therefore, we examined Symbiodiniaceae clades hosted by *A. tenuis* to see whether the clade differs among the five color morphs.

RNA-Seq analyses of the internal transcribed spacer 2 (ITS2) sequence showed that all color morphs hosted only Clade C (*Cladocopium*) zoothanthellae (Fig. 5). No other clades were detected. Among them, group C1 was most abundant, and C50 was next, with smaller numbers of various other Clade-C subgroups (Fig. 5). NB and GB morphs hosted more C3k, whereas P morph hosted more C1i compared with other color morphs, although these differences were not discrete. These symbiont profiles did not change during the study period from late June to early September 2017. *Acropora tenuis* appeared to incorporate higher numbers of C1 and C50 zoothanthella in August and September than in June and July, and this tendency was pronounced in GO color morphs. Nonetheless, these results indicate that shifts in symbiont clades are not the main cause of color differences and/or thermal-resistance of the color morphs, although their levels of photosynthetic activity are likely involved in bleaching of NG and P morphs (Fig. 4).

**Figure 5.**
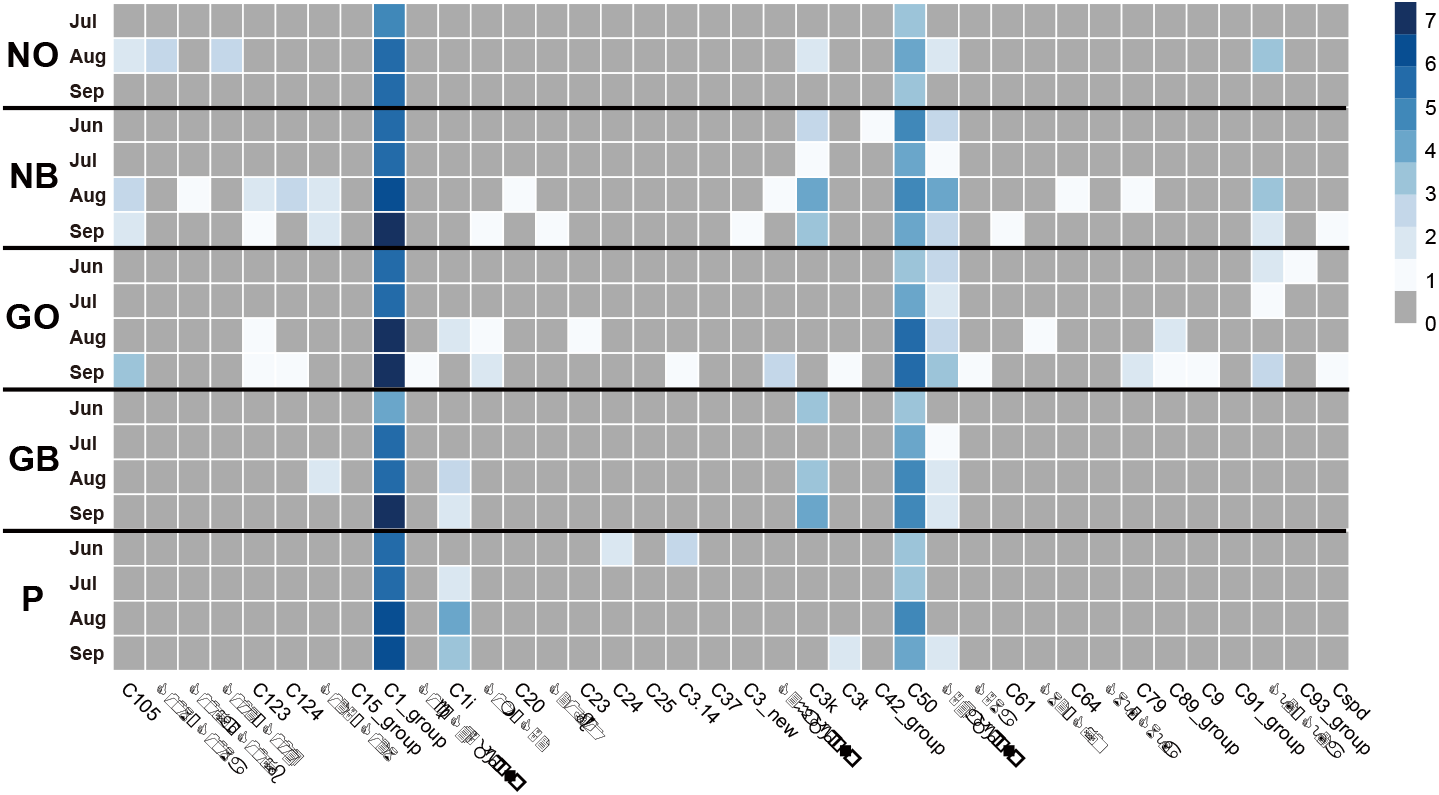
Symbiotic zoothanthellae hosted by five color morphs of *Acropora tenuis*. Zoothanthella clades were identified based upon ITS-2 sequences, determined by RNA-seq analysis. Numbers of sequencing reads are shown by color density (right upper column) as log2 (number of valid alignments +1). All sub-populations hosted only C lade C zoothanthellae (*Cladocopium*), numbers of which did not change during the summer examined.

### Genes for fluorescent proteins

Vivid coloration of corals has been attributed to fluorescent proteins, including green (GFP), red (RFP) and cyan (CFP) proteins, and non-fluorescent blue/purple chromoprotein (ChrP) (e.g., Dove *et al*. 2001; Kelmanson *et al*. 2003; Salih *et al*. 2010; Smith *et al*. 2013). Since genome decoding of *A. digitifera* (Shinzato *et al*. 2011), it has been shown that *A. digitifera* contains five GFP genes, one RFP, one CFP, and three genes for ChrP (Shinzato *et al*. 2012). The approximately 407-Mbp genome of *Acropora tenuis* has also been decoded (Shinzato *et al*. 2020). BLAST searches employing *A. digitifera* fluorescent-protein gene sequences as queries demonstrated the presence in the *A. tenuis* genome of five genes for GFP (Gene ID: s0077.g62, s0297.g27, s0297.g28, s0297.g29, and s0366.g7) (shown by green in Fig. 6a), two genes for CFP (s0010.g7 and s0025.g63) (black in Fig. 6b), three genes for RFP (s0024.g131, s0024g134 and s0217.g35) (red in Fig. 6c), and seven genes for ChrP (s0096.g1, s0096.g3, s0096.g4, s0152.g9, s0152.g11, s0152.g15, and s0182.g24) (purple in Fig. 6d), respectively (https://marinegenomics.oist.jp/gallery/gallery/index.). Molecular phylogeny supports orthologous relationships of the *Acropora tenuis* genes with genes of fluorescent proteins and non-fluorescent chromoproteins of other corals (Fig. S3). Transcriptomes corresponding to these genes have been identified during genome decoding (Shinzato *et al*. 2020), and were used to determine their expression profiles (see below).

**Figure 6.**
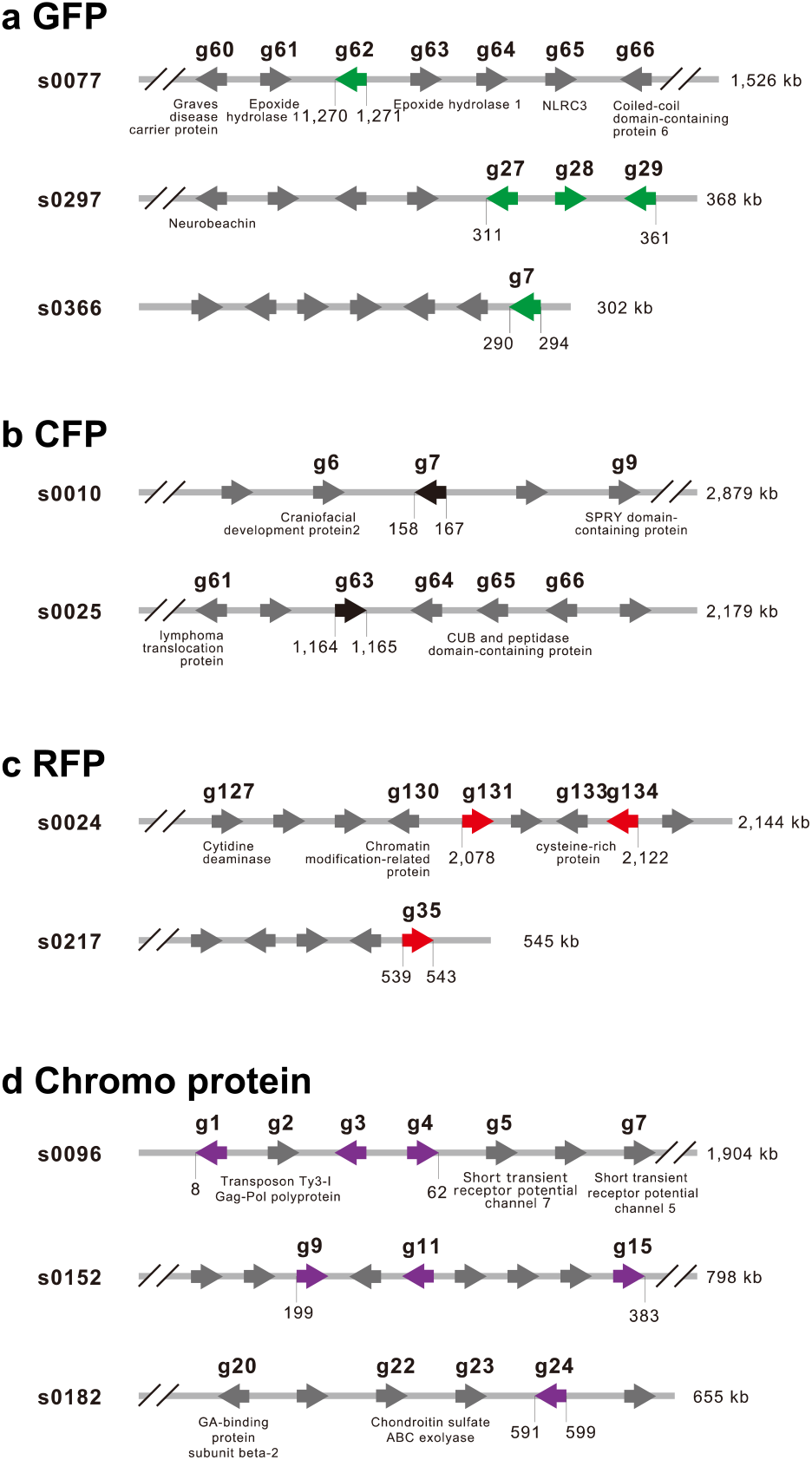
Genes for fluorescent proteins in the *Acropora tenuis* genome. The A. *tenuis* genome contains 13 fluorescent protein-related genes. (a) Three scaffolds, s0077, s0297 and s0366, contain one, three, and one gene for green fluorescent proteins (GFP; shown in green). Other genes (grey) in scaffolds are named if the gene was annotated. Arrowheads indicate the transcription direction. (b) Two scaffolds, s0010 and s0025, contain one gene each for cyan fluorescent proteins (CFP; shown in black). (c) Two scaffolds, s0024 and s0217, contain two and one gene, respectively, for red fluorescent proteins (RFP; shown in red). (d) Three scaffolds, s0096, s0152 and s0182, contain three, three, and one gene for chromoproteins (shown in purple). Molecular phylogeny of these genes is shown in Supplementary Fig. S3.

### Expression profiles of fluorescent protein genes

Ten colonies of four color-morphs (three NB, two GB, three GO and two P) were cultured in an outdoor pool at the Sea Seed Aquarium from late June to early September 2017. We sampled specimens at three time points to examine expression profiles of fluorescent protein genes in each of the ten colonies. Three color morphs (N, G and P) showed different expression profiles of fluorescent protein-related genes during the summer (Fig. 7). First, the two CFP genes and three RFP genes showed very low expression levels with almost no detectable changes during the summer (Fig. 7). This indicates that genes for the four fluorescence-related proteins do not necessarily show similar responses to environmental changes, including surface seawater temperature rise, and that genes for CFP and RFP are unlikely to be regulated by surface seawater temperatures. On the other hand, GFP and ChrP genes did show detectable changes in expression profiles.

**Figure 7.**
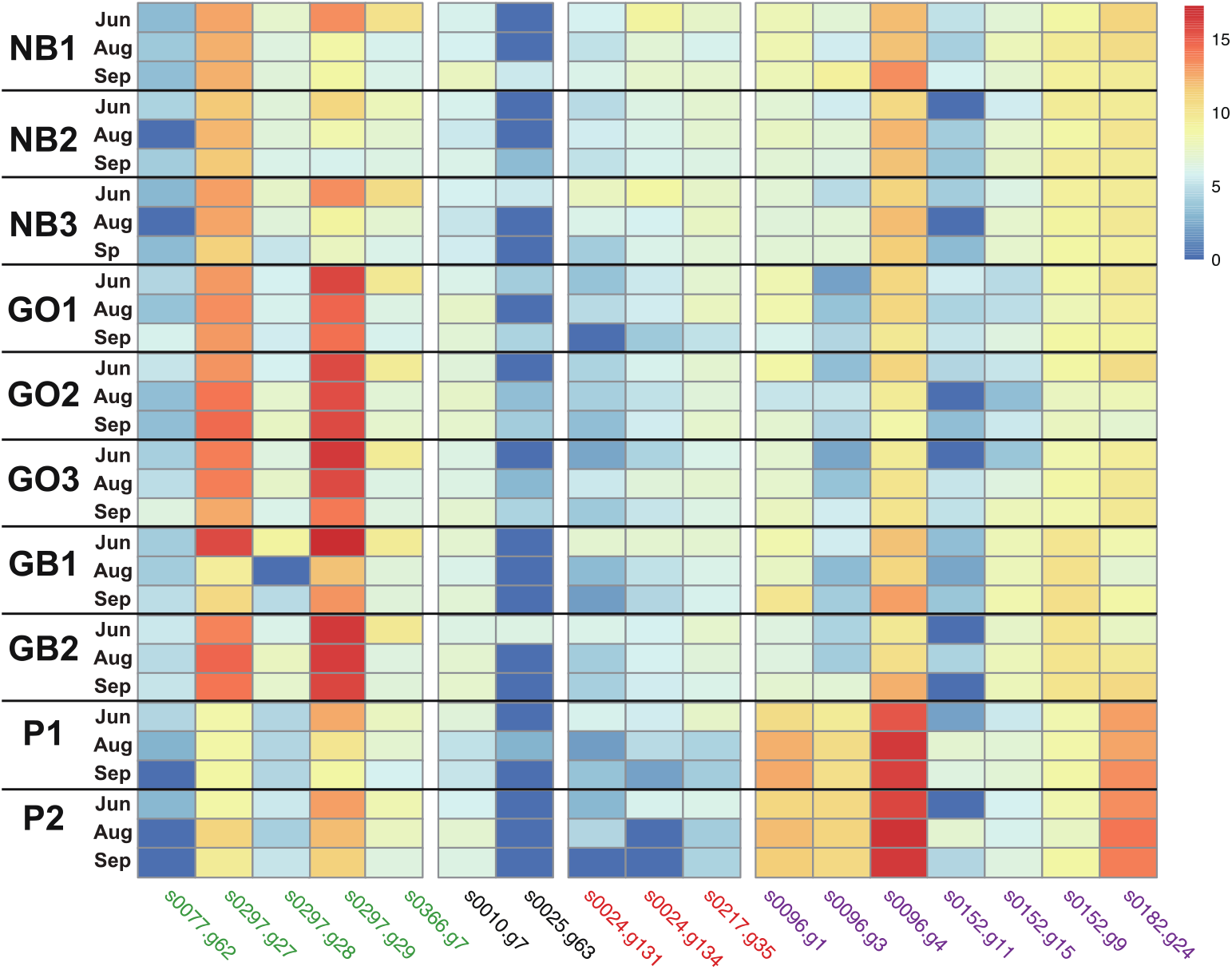
An expression profile of genes for fluorescent proteins in four *Acropora tenuis* color morphs during the summer of 2017. Expression levels of four GFP genes (green), two CFP genes (black), three RFP (red) and seven chromoprotein genes (purple) are shown with their densities (right upper column) as log2 (number of valid alignments +1). Gene IDs are identical to those in Fig. 6. NB1, 2, and 3 are three genotypes of the NB morph, GB1 and 2 are two genotypes of the GB morph, GO1, 2, and 3 are three genotypes of the GO morph, and P1 and 2 are two genotypes of the P morph.

Two GFP genes (s0297.g27 and s0297.g29) showed distinctive changes in expression profile, while expression levels of the other three genes (s0077.g62, s0297.g28 and s0366.g7) remained low, with little change (Fig. 7). This indicates that even among genes that encode the same fluorescent protein and that are located in close proximity on the same chromosome (Fig. 6), their responses to SST are not always the same, suggesting complicated individual gene regulation in response to environmental stresses. In addition, s0297.g27 and s0297.g29 exhibited an interesting shared expression profile during the study period. First, the expression level of s0297.g29 was distinctively higher in GB and GO than in NB and P (Fig. 7). In all three wild-type NB1, 2 and 3 genotypes, s0297.g29 showed moderate levels of expression in June, but diminished levels with rising temperatures in August and September. A similar expression profile of s0297.g29 was detected in GB1, GO1, GO3, and P1. However, gene expression was distinctively different between N and G. The expression level was much higher in GB and GO than in NB. In addition, the higher level of s0297.g29 expression in GB2 and GO2 was maintained throughout the summer (Fig. 7). Although less obvious, a similar expression profile was observed in s0297.g27 (Fig. 7).

The purple morphs exhibited another expression profile of these genes, especially in relation to genes for non-fluorescent blue/purple chromoprotein (ChrP). Of seven ChrP genes, s0096.g1, s0096.g3, s0096.g4 and s0182.g24 exhibited an interesting expression profile (Fig. 7). Although the level of the three other genes, s0152.g9, s0152.g11 and s0152.g15 (localized in the same scaffold) was comparatively low (Fig. 7), the former four ChrP genes were expressed at higher levels in P1 and P2 than in NB, GB, and GO (Fig. 7). Although expression level differed among the four genes, this pattern was shared by all four color morphs and relative levels of the four genes were maintained during the summer as surface seawater temperatures rose (Fig. S2a). s0096.g4 showed significantly higher expression in both P1 and P2 as September approached (Fig. 7). This tendency was also seen in s0182.g24. These results indicate that in contrast to the role of GFP in N and G colonies, ChrP is very likely involved in coloration of P colonies and has potential to resist surface seawater temperature rise.

## DISCUSSION

There are at least three color morphs of *Acropora tenuis* along the Okinawa coast, brown, green and purple (Nishihira and Veron 1995) (Fig. 1). They exhibit different profiles of fluorescence in the axial polyps when excited at 405 nm (Fig. 2). We expect that differences in fluorescence are associated with coloration of this coral, although detailed molecular, biochemical, and biophysical mechanisms underlying these differences should be addressed in future studies. The color morphs are stable as far as we have observed for a decade, since color morphs show no seasonal alternation (e.g., Paley 2014), although the brightness of color changes. Color is asexually heritable. Although it remains to be determined whether it is also sexually heritable, color polymorphism is likely caused by genetic / genomic variation.

Field studies in 2017 showed that N morphs underwent extensive bleaching (Fig. 3a, b), and 10∼15% of NB morphs eventually died (Fig. 3b). Bleaching occurred in P morphs in late August, but many of them recovered in subsequent months (Fig. 3e). In contrast, G morphs did not show bleaching (Fig. 3c, d). Therefore, it is evident that sensitivity to thermal stress differs among the three color morphs. N is the most sensitive, P is moderate, and G is resistant. Along the Okinawa coast, the N color morph is most abundant (as seen in Fig. 3 and Fig. S1), while G and P are not as common. This suggests that the N morph is probably ancestral, while the G and P color morphs are more recent. If this feature is heritable, it is tempting to speculate that in Okinawa, *A. tenuis* has acquired the capacity to resist thermal stresses by developing new color morphs. In other words, polyp color polymorphism may be a strategy to survive severe summer environmental stresses, provided that the current rate of temperature change does not overwhelm the coral’s capacity to adapt.

Bleaching responses are complicated, varying across colonies, taxa, and events (e.g. van Woesik *et al*. 2011). Scleractinian corals form obligate endosymbioses with photosynthetic dinoflagellates or zoothanthellae, and host-symbiont interactions contribute to differential bleaching susceptibility (Enriquez *et al*. 2005; Hawkins *et al*. 2014; Wooldridge 2014). Since bleaching is usually caused by escape and/or death of symbionts, declining photosynthetic activity of zoothanthellae may portend bleaching. Indeed, decreased photosynthetic activity was detected in June in NB and P morphs (Fig. 4). In contrast, in July, GO and GB morphs maintained photosynthetic activity comparable to that in other months. This suggests that G color morphs provide a suitable physiological environment for symbionts to remain in the host despite the high thermal stress. Genetic and molecular mechanisms involved in this relationship are intriguing, and should be addressed in future studies.

The zoothanthella family Symbiodiniaceae has recently been reorganized into nine clades or genera (A-I), according to new analyses of combinatorial data (LaJeunesse *et al*. 2018). It has been suggested that in general, corals hosting Clade D (*Durusdinium*) are more heat-resistant than those hosting Clade C (*Cladocopium*) or that specific Symbiodinium phylotypes, such as D1, D1-4, C15 and A3 are exceptionally thermotolerant, while others (e.g. C3, C7, B17, A13) are thermosensitive (e.g. Silverstein *et al*. 2011; Tonk *et al*. 2014). Therefore, differing capacities of *A. tenuis* color morphs to resist higher surface seawater temperatures may be due to different zoothanthella hosted by the three color morphs. However, this is not the case in Okinawa *A. tenuis*, because all three host a very similar repertoires of Clade C *Symbiodinium*, especially C1 and C5 types (Fig. 5).

*Acropora tenuis* is useful as an experimental system because its ∼400-Mb genome has been decoded, and is thought to contain 22,802 protein-coding genes (Shinzato *et al*. 2020). Moreover, approximately 95% of the gene models are confirmed with corresponding mRNAs. Therefore, we were able to investigate not only the genes for GFP, CFP and RFP and ChrP, but also their expression profiles in colonies with different color morphs during the summer of 2017. We found that all three morphs possess an identical array of fluorescence-related genes and that expression of these genes was confirmed in all morphs using RNA-seq. This indicates that color polymorphism is not caused by mutations in the genes themselves. Instead, molecular and genic mechanisms that control transcriptional activity or quantitative regulation over gene products may produce the three color morphs. Genetic and genic mechanisms underlying fluorescent protein-mediated color polymorphisms and adaptation potential to variable environmental conditions have been studied in *Acropora millepora* as an experimental system (D’Angelo *et al*. 2008; Smith *et al*. 2013; Gittins *et al*. 2015). In this species, an RFP gene, *amiFP597*, is essential for color polymorphism. The gene, *amiFP597*, is multicopy with a particular promoter type in the genome, and the number of gene copies is strongly correlated with the level of gene expression. Higher levels of gene expression are found in more intensely red morphs, which show higher resistance to strong insolation (Gittins *et al*. 2015). This suggests the presence of variable genetic mechanisms in color polymorphisms, depending on coral species.

Interestingly, the P morph exhibits a gene expression profile different from those of the N and G morphs. Specifically, the P morph shows higher ChrP gene expression, especially s0096.g4 (Fig. 7). In addition, the N and G morphs differ in expression levels of GFP genes, especially s0297.g29 and s02897.g27 (Fig. 7). Higher gene expression was evident in the G morph compared to the N morph (Fig. 7). Because the G morph is the most resistant to high SSTs, it is likely that stress resistance varies with different expression levels of GFP genes. We expect that different control mechanisms of GFP gene expression in N and G morphs explain their different thermal tolerance. We will address this question in future studies.

Coral bleaching is caused by multiple, complex environmental stresses. Except for outbreaks of crown-of-thorns starfish, typhoons, and diseases, the most deleterious influence is surface seawater temperature warming. Strong solar radiation also likely contributes to environmental stress. Seawater in Okinawa Prefecture is very transparent; thus, UV irradiation acts in concert with higher sea temperatures to cause bleaching. Fluorescent proteins and chromoproteins absorb UV radiation, rendering corals and their symbionts more resistant to solar stress (Dove *et al*. 2001; Dove 2004; Banaszak and Lesser 2009; Salih *et al*. 2010). We need to better understand the relationship between solar stress and different expression profiles of genes for GFP and ChrP.

Divers and marine researchers have noticed that some corals, sometimes called “Super corals” (https://phys.org/news/2019-05-super-corals-glimmer-world-dying.html) and/or “super coral reefs” (Dance 2019), are able to survive severe environmental stresses. Nonetheless, almost nothing is known about how such super corals have appeared in the reefs. Our present results regarding different thermal tolerances of three *Acropora tenuis* color morphs may help to explain the resilience of these super corals.

## ACKNOWLEDGEMENTS

This work was supported by Okinawa Prefecture Project of Coral Preservation to KS, DK, HI, AY, KH and NS. The Sumiko Imano Memorial Foundation by Sumio Imano, Haruo Imano and Hiromichi Imano to NS and OIST support to Marine Genomics Unit are much appreciated. YZ and CS are supported by JSPS grants (17K15179 and 19K15902 to YZ, and 17K07949, 17KT0027 and 20H03235 to CS). The DNA Sequencing and Imaging Sections of OIST are gratefully acknowledged. We thank Dr. Steven D. Aird for editing the manuscript.

## Figure Legends

**Supplementary Figure S1.**
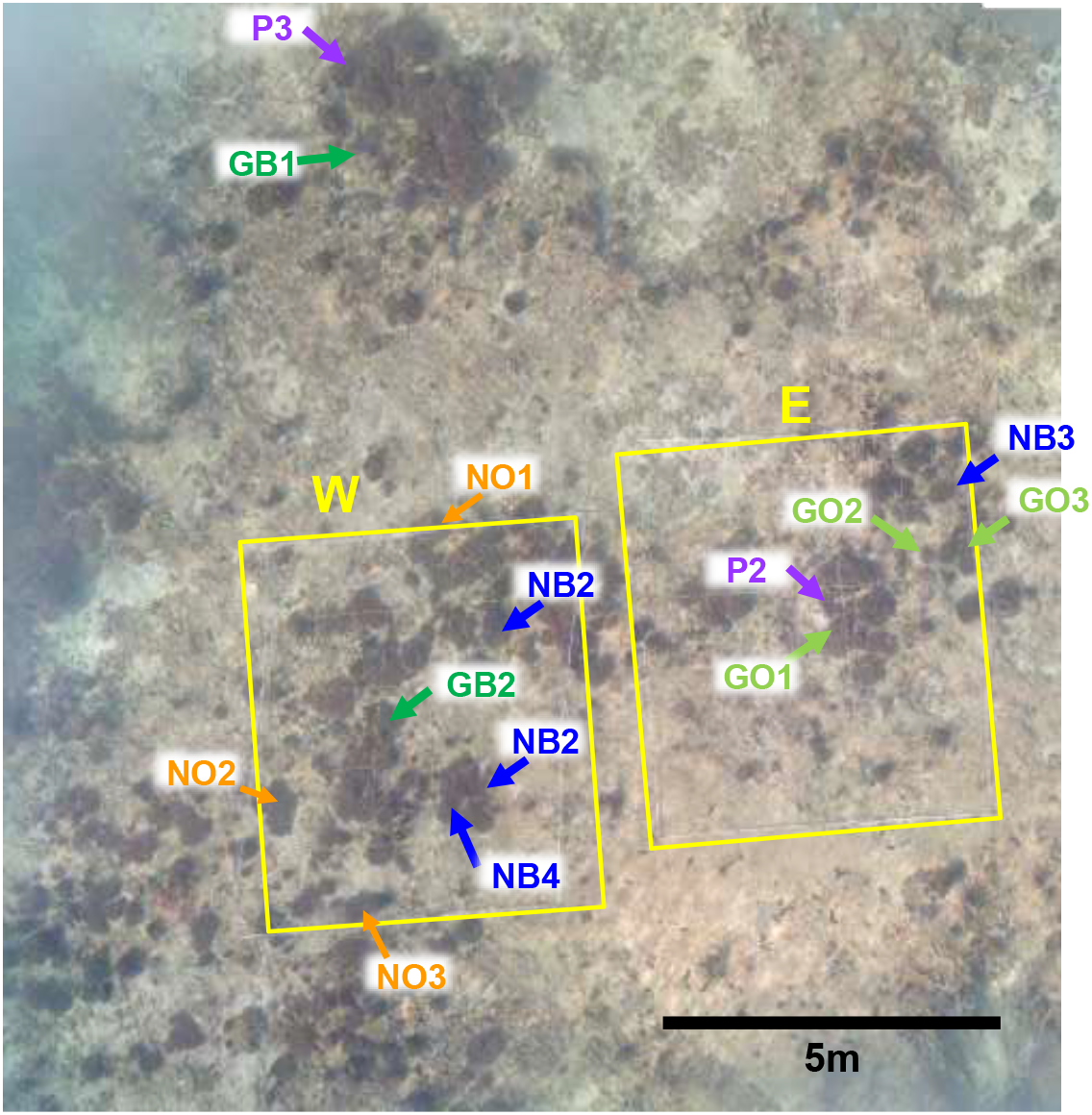
An ortho-image of a sample site of *Acropora tenuis* to monitoring bleaching of the five color morphs in 2017. Two quadrats (2m X 2m), enclosed with yellow, contain seven (left) and five morphs with different colors, which were used for PAM measurement of photosynthetic activity. Bleaching was observed on all morphs in the quadrats.

**Supplementary Figure S2.**
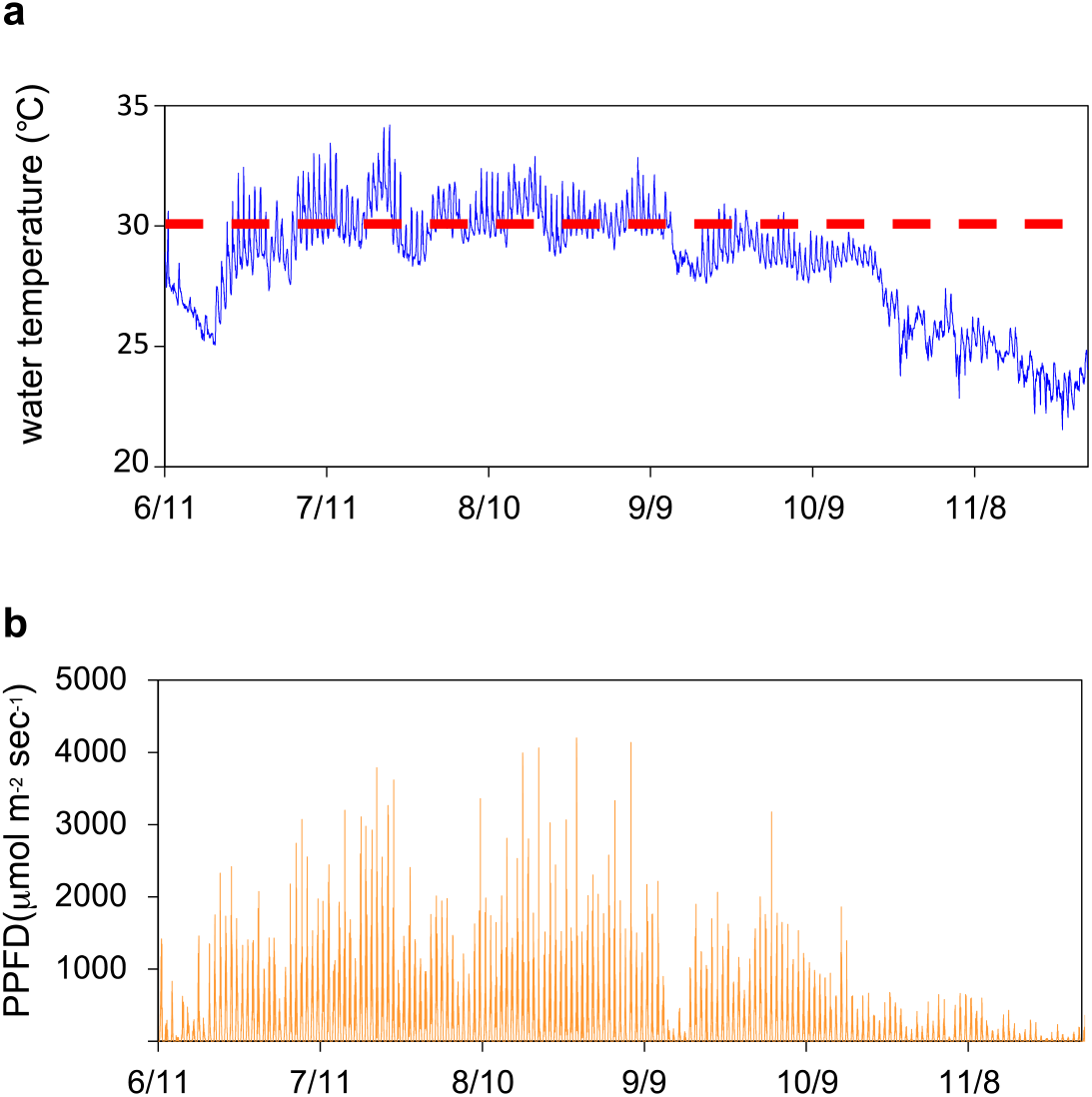
Changes in surface seawater temperature (SST) (a) and photosynthetic photon flux density (b) during the summer and autumn of 2017. Both were automatically measured. June 11, July 11, August 10, September 9, October 9 and November 8 indicate dates of coral bleaching observations. SSTs >30°C continued from early July to early September. The dotted red line indicates 30°C.

**Supplementary Figure S3.**
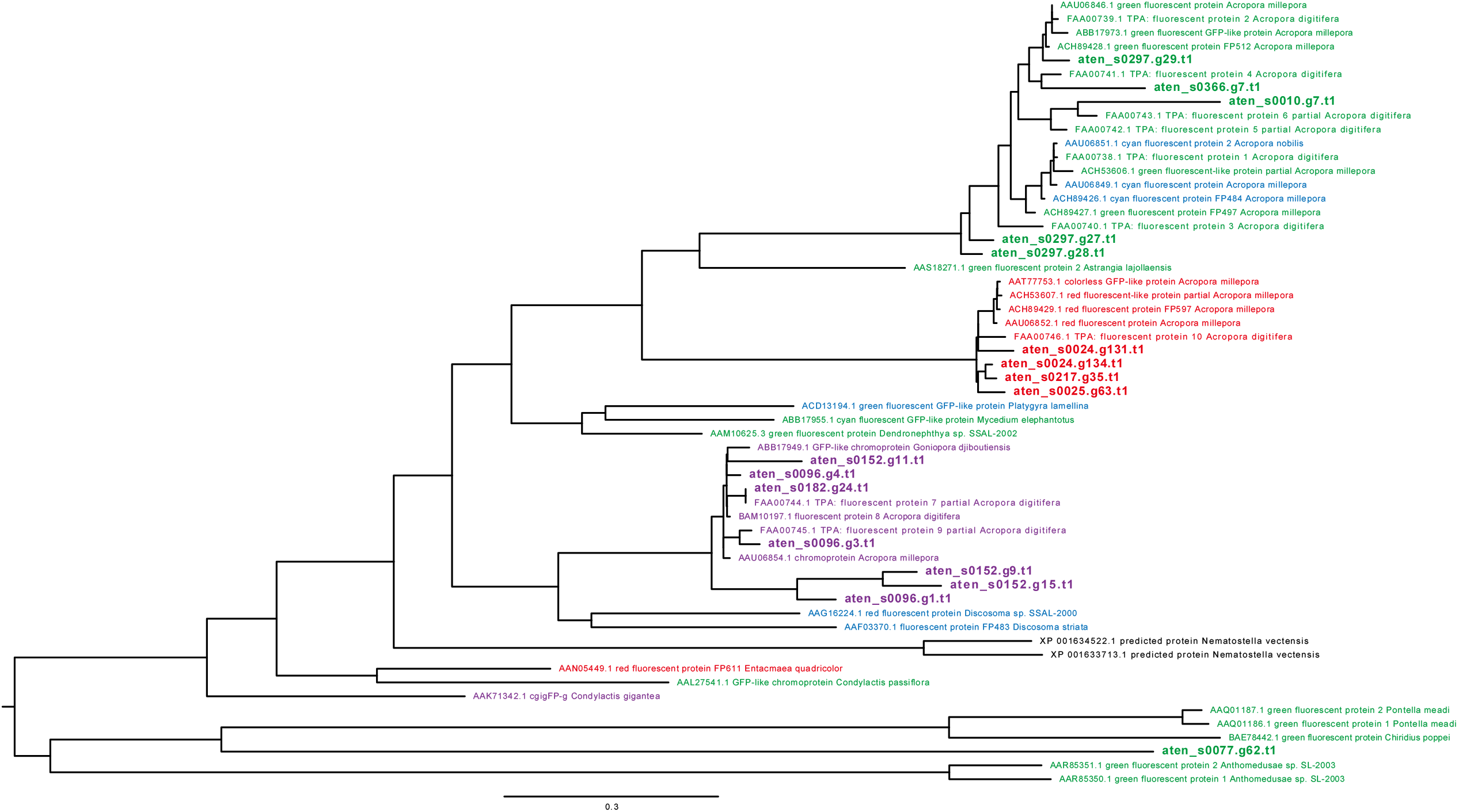
Molecular phylogeny of coral fluorescent proteins of *Acropora tenuis*. Phylogenetic relationships of four GFPs (green bold letters), two RFPs (red bold letters) and seven chromoproteins (purple bold letters) are shown. CFPs are not included due to their extreme divergence from the others. The tree was constructed using the NJ method and bootstrap values (% of 1,000 replicates) are shown for major branches. The scale bar indicates a 0.5% substitution rate.

